# Autistic traits, resting-state connectivity and absolute pitch in professional musicians: shared and distinct neural features

**DOI:** 10.1101/456913

**Authors:** T. Wenhart, R.A.I. Bethlehem, S. Baron-Cohen, E. Altenmüller

## Abstract

**Background:** Recent studies indicate increased autistic traits in musicians with absolute pitch and a higher incidence of absolute pitch in people with autism. Theoretical accounts connect both of these with shared neural principles of local hyper- and global hypoconnectivity, enhanced perceptual functioning and a detail-focused cognitive style. This is the first study to investigate absolute pitch proficiency, autistic traits and brain correlates in the same study.

**Sample and Methods:** Graph theoretical analysis was conducted on resting state (eyes closed and eyes open) EEG connectivity (wPLI, weighted Phase Lag Index) matrices obtained from 31 absolute pitch (AP) and 33 relative pitch (RP) professional musicians. Small Worldness, Global Clustering Coefficient and Average Path length were related to autistic traits, passive (tone identification) and active (pitch adjustment) absolute pitch proficiency and onset of musical training using Welch-two-sample-tests, correlations and general linear models.

**Results:** Analyses revealed increased Path length (delta 2-4 Hz), reduced Clustering (beta 13-18 Hz), reduced Small-Worldness (gamma 30-60 Hz) and increased autistic traits for AP compared to RP. Only Clustering values (beta 13-18 Hz) were predicted by both AP proficiency and autistic traits. Post-hoc single connection permutation tests among raw wPLI matrices in the beta band (13-18 Hz) revealed widely reduced interhemispheric connectivity between bilateral auditory related electrode positions along with higher connectivity between F7-F8 and F8-P9 for AP. Pitch naming ability and Pitch adjustment ability were predicted by Path length, Clustering, autistic traits and onset of musical training (for pitch adjustment) explaining 44% respectively 38% of variance.

**Conclusions:** Results show both shared and distinct neural features between AP and autistic traits. Differences in the beta range were associated with higher autistic traits in the same population. In general, AP musicians exhibit a widely underconnected brain with reduced functional integration and reduced small-world-property during resting state. This might be partly related to autism-specific brain connectivity, while differences in Path length and Small-Worldness reflect other ability-specific influences. This is further evidence for different pathways in the acquisition and development of absolute pitch, likely influenced by both genetic and environmental factors and their interaction.

## Background

Autism spectrum disorders or conditions (henceforth ‘autism’) are more common in people with mathematical [1], visuo-spatial [2], musical [3] or ‘savant’ abilities [4], e.g. rapid mental mathematical calculation [5, 6], calendar calculation [7], or extreme memory [8, 9]. Autism, a set of neurodevelopmental condition, are characterized by social and communication difficulties, alongside unsually repetitive behaviors and unusually narrow interests [10], sensory hypersensitivity, and difficulties in adjusting to unexpected change (DSM-5, APA 2013).

Absolute pitch (AP), the ability to name or produce a musical tone without the use of a reference tone [11] is a common special ability in professional musicians with an incidence of up to 7-25% [12–14] but less than 1% [15] in the general population. AP is an excellent model for the investigation of a joint influence of genetic and environmental factors on the brain and on human cognitive abilities [16]. An influence of age of onset of musical training [17–19], ethnicity [12, 14, 19], and type of musical education (label to fixed pitch vs. label to interval, unfixed to pitch) techniques [12]) suggest environmental aspects in the acquisition of AP. In contrast, AP often clusters in families, genetically overlaps with other familial aggregated abilities (e.g. synesthesia [20]) and has a higher incidence in autistic people [3, 7, 21–25] and in Williams-syndrome [26], both strongly genetic conditions [27–34]. Finally, a sensitive or critical period before the age of seven is considered due to the importance of early onset of musical training [14, 16, 17, 35–38] Recently, two studies have given evidence for heightened autistic traits in musicians with AP [39, 40]. Both AP and autism are associated with similarly altered brain connectivity in terms of the relation between hyper- and hypo-connectivity [36, 41–50]. The theory of veridical mapping [7] tries to explain absolute pitch, synesthesia and other abilities like hyperlexia, frequently seen in autistic people or in savant syndrome, with the neurocognitive mechanism of associating homologues structures of two perceptual or cognitive structures (veridical mapping). According to this framework, enhanced low level perception [51, 52] and an increased ability to detect patterns (‘systemizing’ [53]) is associated with regional hyper- as well as global hypo-connectivity in absolute pitch [41, 43, 54–59] and autism [42, 44, 46, 60]. It is also noteworthy that autism and abilities like absolute pitch share excellent attention to detail [35, 61] and a shift in the direction of higher segregation with reduced integration in the brain [61]. Investigating disconnection syndromes or integration deficit disorders, as well as phenomena with similar brain network characteristics, may therefore provide insights into the variability of brain network structure and function and its relation to perception, cognition and behaviour.

The present study tests if and to what extent AP and autistic traits share the same neurophysiological network connectivity. To our knowledge, this study is the first to investigate (1) the relation of pitch adjustment ability (active absolute pitch; in contrast to (passive) pitch identification) and brain as well as behavioral correlates, (2) the relation of AP ability, autistic traits and functional brain connectivity within one study, and (3) graph theoretical network parameters in AP during resting state electroencephalography. We use graph theoretical analysis [62, 63] of resting state EEG data to estimate differences in global network structure of the brain. We analyzed three graph theoretical network parameters reflecting segregation (Average Clustering Coefficient) and integration (Average Shortest Path Length) and so called Small-Worldness (a combination of Clustering and Path length) [62, 63]. To our knowledge this is also the first study investigating global i.e. average connectivity parameters over the whole brain between AP and RP (relative pitch) musicians, while prior studies [41, 43] have focused on parameters for single regions (e.g. degree, single node clustering, and single node characteristic path length). We expected higher autistic traits, higher Path length (reduced integration) and lower Clustering (underconnectivity) for AP and an interrelation among those variables. Further, we expected these differences to specifically occur in low (delta, theta) vs. high frequency (beta) ranges for integration vs. segregation, respectively.

## Methods

### Participants

Thirty-one AP musicians (16 female) and 33 RP musicians (15 female) participated in the study. One male RP participants had to be excluded from EEG analysis because of missing EEG-data. Participants were recruited via an online survey using UNIPARK software (https://www.unipark.com/) and primarily were students or professional musicians at the University for Music, Drama and Media, Hanover. Four AP and two RP were amateur musicians. An online pitch identification screening (PIS) consisting of 36 categorical, equal-tempered sine waves in the range of three octaves between C4 (261.63 Hz) and B6 (1975.5 Hz) was used to allocate the participants to the groups (AP: >12/36 tones named correctly, else RP). Four AP were non-native German speakers and had the choice between a German and an English version of the experiments. One AP reported taking Mirtazapine. None of the participants reported any history of severe psychiatric or neurological conditions. The AP group consisted of 15 pianists, 9 string players, 3 woodwind instruments, two singers and 2 brass players; the RP group consisted of 13 pianists, 4 string players, 6 woodwind instruments, 3 bassists/guitarists/accordionists, 3 singers, one drummer and 3 brass players. Handedness was assessed by Edinburgh Handedness Inventory [64]; one AP was left handed, all other AP were consistently right handed, three RP were left-handed, two RP were ambidextrous. This study was approved by the local Ethics Committee at the Medical University Hannover. All participants gave written consent.

### Setting

The study was divided into three parts: the online survey and two appointments in the lab at the Institute for Music Physiology and Musicians Medicine of the University for Music, Drama and Media, Hannover. While the online survey was used for the pitch identification screening and diagnostic as well as demographic questionnaires (see below), general intelligence, musical ability, pitch adjustment ability and resting state EEG were assessed in the lab (see Table 1). Four further experiments were conducted within the same two sessions at the lab and are reported elsewhere. Raven’s Standard Progressive Matrices [65] and “Zahlenverbindungstest“ (ZVT, [66]) were used to assess general nonverbal intelligence and information processing speed, respectively. Musical ability and musical experience were controlled for with the use of AMMA (Advanced Measures of Music Audiation, [67]), Musical-Sophistication Index (GOLD-MSI, [68]) and estimated total hours of musical training within life span (house intern online questionnaire).

**Table 1.** Participants’ characteristics

|  | AP (n=31) |  |  | RP (n=33) |  |  | t-test |
| --- | --- | --- | --- | --- | --- | --- | --- |
|  | Mean | SD | Range | Mean | SD | Range |  |
| age | 25.13 | 9.2 | 17-58 | 24.0 | 7.02 | 17-57 | t(56.1)= -0.549;<br>p = 0.585 |
| SPM-IQ | 110.4 | 16.4 | 73-132.25 | 114.41 | 13.14 | 86.5-134.5 | t(57.5)= 1.073;<br>p = 0.288 |
| ZVT-IQ | 120.76 | 13.14 | 101.5-145 | 120.61 | 13.69 | 97-143.5 | t(61.9)= -0.045;<br>p = 0.964 |
| hours main instrument | 11961.4 | 9212 | 1642.5-39785 | 13735.61 | 17125.89 | 1606-77617.25 | t(49.7)= 0.520;<br>p = 0.605 |
| AMMA | 64.74 | 6.26 | 53-78 | 63.244 | 7.03 | 46-76 | t(61.8)= -0.90;<br>p = 0.370 |
| MSI | 208.65 | 17.59 | 161-234 | 210.79 | 15.12 | 185-246 | t(59.3)= 0.521;<br>p = 0.604 |
| **PIS** | 28.5 | 6.03 | 15-36 | 5.30 | 4.33 | 0-21* | **t(52.2)= -17.37;**<br>**p < 2.2e-16** |

Age, nonverbal IQ (SPM), information processing capacity (ZVT), musical training (total hours during life span on main instrument), musicality (AMMA; MSI) and online pitch identification screening (PIS) for each group; \* two RP reported not having absolute pitch but reached a screening score of 13 respectively 21. Because of this and their weak performance in the pitch adjustment test, the subjects were assigned to the RP group; Significant group differences highlighted in bold.

### Experiments and material

#### Pitch Adjustment Test (PAT)

Absolute pitch ability was measured by using two different absolute pitch tests: The pitch identification screening (PIS) during the online survey mentioned above, and a pitch adjustment test (PAT) based on Dohn et al. [69]. Participants were given a maximum of 15 seconds to adjust the frequency of a sine wave with random start frequency (220 −880 Hz, 1Hz steps) and told to try to hit the target note (letter presented central on PC screen, e.g. “F# / Gb”) as precisely as possible without the use of any kind of reference. Online pitch modulation was programmed according to Dohn et al. [69] and provided by turning a USB-Controller (Griffin PowerMate NA16029, Griffin Technology, 6001 Oak Canyon, Irvine, CA, USA). Resolution of the Power Mate was set to 10 cents vs. 1 cent (if pressed during turn of the wheel) for individual choice between rough and fine tuning. To confirm their answer, participants were instructed to press a button on a Cedrus Response Pad (Response Pad RB-844, Cedrus Corporation, San Pedro, CA 90734, USA) to automatically proceed with the next trial. If no button was hit, the final frequency after 15 seconds was taken. In both cases, the Inter Trial Interval (ITI) was set to 3000 ms. The total test consisted of 108 target notes, presented in semi-random order in 3 Blocks of 36 notes each (3*12 different notes per block) with individual breaks between the blocks. The final or chosen frequencies of each participant were compared to the nearest target tone (< 6 semitones/600cent), as participants were allowed to choose their octave of preference. EEG was measured during the PAT but will be reported elsewhere. For each participant, mean absolute derivation (MAD (1),[69]) from target tone
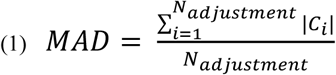

is calculated as the mean of the average absolute deviations c_i_ (2) of the final frequencies to the target tone (referenced to a 440 Hz equal tempered tuning).

MAD reflects the pitch adjustment accuracy of the participants. The consistency of the pitch adjustments, possibly reflecting the tuning of the pitch template[69], is then estimated by taking the standard deviation of the absolute deviations (2).
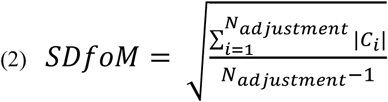

For regression analyses (see below), we performed a z-standardization of the MAD (Z_MAD,(3)) and SDfoM (Z_SDfoM, (4)) values relative to the mean and sd of the non-AP-group, as originally proposed by Dohn et al. [69].
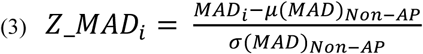

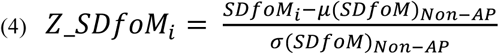

#### Autistic Traits

Autism traits were assessed during the online survey using a standardized Adult Autism Spectrum Quotient (AQ, [70]; German version by C.M. Freiburg, available online: https://www.autismresearchcentre.com/arc_tests). It consists of 50 items within five subscales (attention to detail, attention switching, imagination, social skills and communication). One point is given for each item with a mildly or strongly agreement with the autistic-like symptoms (half the items were negatively poled. The maximum AQ-Score therefore is 50).

#### EEG Resting State

EEG resting state data was acquired immediately before the PAT at the beginning of the experimental session using 28 scalp electrodes (sintered silver/silver chloride; Fp1, Fp2, F3, F4,FC3, FC4, C3, C4, CP3, CP4, P3, P4, F7, F8, FT7, FT8, T7, T8, TP7, TP8, P7, P8, O1, O2, Oz, Fz,Cz, Pz) placed according to the international extended 10-20 System with an electrode cap by EASYCAP (EASYCAP GmbH, Herrsching, Germany; http://www.easycap.de). A 32-channel SynAmps amplifier (Compumedics Neuroscan, Inc., Charlotte, NC, USA) and the software Scan 4.3 (Compumedics Neuroscan) were used to record the data. The remaining 2 bipolar channels were used for vertical and horizontal electro-oculogram with electrodes placed above and below the right eye and approximately 1cm outside of the outer canthus of each eye, respectively. Two further electrodes were placed on the left and right mastoids as a linked reference. The ground electrode was placed between the eyebrows directly above the nasion on the forehead. Abralyth 2000 abrasive chloride-free electrolyte gel (EASYCAP GmbH, Herrsching, Germany; http://www.easycap.de) was used to keep impedances below 5kΩ. Participants were seated in a comfortable chair in front of a PC screen and were instructed to let their mind wander around while looking at a fixation cross (eyes open resting state, EO) or keeping their eyes closed (eyes closed resting state, EC) for 5 minutes each. Start (button press) and end of the resting state period where programmed within PsychoPy [71] by sending triggers via a parallel port to the EEG-system. A sampling rate of 1000 Hz was used combined with an online-bandpass filter between 0.5-100Hz and a Notch-filter at 50 Hz. EEG was recorded in AC (alternating current)-mode and with a gain of 1000.

### EEG Preprocessing and Analysis

#### Preprocessing

All preprocessing steps were conducted using MATLAB (MATLAB Release 2014a, MathWorks, Inc., Natick, Massachusetts, United States) using the toolboxes eeglab [72] and fieldtrip [73]. EEG raw data was first re-sampled to 512 Hz sampling rate and bandpass filtered to 1-100 Hz. Artefact removal was administered using both, raw data inspection of continuous data and independent component analysis (ICA, algorithm: binica) within eeglab for each participant’s data individually. ICA-components containing vertical or horizontal eye movements, blinking, heartbeat, muscular activity or other artefacts were removed from the data by inverse ICA. After that, segments still containing the above mentioned artefacts were removed manually. Defective or highly noisy electrodes were interpolated using spherical interpolation [74] implemented in eeglab (5 participants, 1-2 electrodes each). All statistical analyses were repeated under exclusion of participants with interpolated electrodes as well as non-native German speakers and the participant which reported to take Mirtazapine. Direction and significance of effects was not affected by the exclusions, therefore all participants were included into the final analyses. Afterwards, the artefact clean data was exported to fieldtrip for connectivity and network analysis (next steps).

#### Connectivity – weighted Phase Lag Index (wPLI)

Calculation of functional connectivity was done using MATLAB scripts (see: https://github.com/rb643/fieldtrip_restingState/blob/master/rb_EEG_Conn.m). First, 4s-epochs (non-overlapping) were extracted from the artefact-clean data. Second, multi-taper Morlet fast Fourier transformation was used to extract frequency bands (delta: 2-4 Hz, theta: 4-7 Hz, alpha: 7-13 Hz, beta: 13-30 Hz, gamma: 30-60 Hz). For delta and theta single-taper (Hanning window) was used. Contrary, for alpha, beta and gamma multiple tapers (discrete prolate spheroidal sequences, DPSS) were taken. During multi-tapering of alpha, beta and gamma spectral smoothing was applied (+-1,2,4 Hz, respectively). Finally, pairwise connectivity values for each electrode site were calculated per participant and stored in a connectivity matrix for each frequency band separately. Weighted phase lag index [75] was chosen as connectivity measure, as phase based connectivity measures compared to coherence and phase synchronization measures are less sensitive to volume conduction in the brain [76, 77] cited by [78]), i .e. spurious connectivity between two regions of interest caused by a common source of activity or a common reference [79] and usually leads to connectivity values with phase lags of zero or pi (if the two sites are on opposite sides of the dipole) [80]. PLI (5)
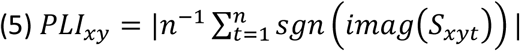

is an index that quantifies the asymmetry of the distribution of instantaneous phase-differences ΔΦ between the signals x and y, by averaging the sign (sgn) of the imaginary components (imag) of the cross-spectrum (S_xyt_) at timepoint t[80].The distribution is centered around 0 mod pi, therefore an asymmetric distribution shows non-zero phase lag. Stam et al. [79] argue, that a non-zero phase lag cannot be caused by volume conduction or a common reference, as the latter work instantaneously. PLI takes values between 0 and 1, where 0 indicates no phase coupling (or a coupling with a 0 mod pi phase difference) and 1 indicates a perfect coupling at the phase lag of ΔΦ. Because of the absolute values taken in equation (1) PLI does not give information about which signal is leading [79]. PLI has been shown to be superior in detecting true synchronization and in being less influenced by common source activity and electrode montage systems than phase coherence (PC, [81]), both in computer simulations and on real EEG and MEG data [79]. Furthermore, PLI exhibits a similar amount of long to short distance connections in an investigation of beta-band coupling in Alzheimer data [79, 82] which was shifted towards short-range connections implying volume conduction when using PC [79]. As the aim of this paper is to compare graph theory based network measures that especially quantify segregation versus integration in the brain (see section Network analysis – Graph Theory) the use of PLI is to be preferred to prevent the distortion of the network parameters by volume conduction [65, 66]. The extension of PLI, weighted PLI (wPLI, (6) [75]),
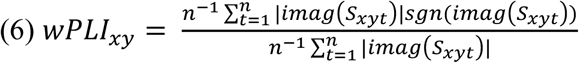

weights the obtained phase leads or lags by the magnitude of the imaginary component (imag) of the cross-spectrum (S_xyt_). This reduces the influence of additional noise sources [75, 80]. Weighted phase-lag-index [75] therefore is an advancement of phase lag index (PLI, [79]) and a suitable measure to detect true connectivity between regions of interest [79], as it ignores zero- and pi-phase-lag.

#### Network analysis - Graph Theory

Graph network analyses were conducted using Brain Connectivity Toolbox (BCT,[85]) in Matlab. Graph theory is a branch of mathematics that deals with the abstract representation of networks as graphs, i.e. a system of n nodes and k edges (connections) between the nodes. Increasingly, network science is being applied to a range of neuroanatomical and -physiological data (e.g. [41, 43, 86–91]) and at different scales of interest (e.g. neurons/populations of neurons, cortical areas, electrode sites; see [62, 85, 92–94] for an overview). In the present study, the pairwise connectivity measures for each frequency band and participant were stored in a 28×28 (channel by channel)-matrix. Therefore, electrode sites are defined as nodes and the wPLI indexes of the electrode pairs within the matrix as edges. This was done in two steps: First, to construct adjacency matrices for graph analyses, minimal spanning tree (MST: van Wijk et al., 2010) was used as the threshold starting point for building binary networks at various densities. The density of a network relates to the fraction of edges present in the network compared to the maximum possible number of edges. MST was chosen to ensure that across participant we were comparing network with similar numbers of nodes (e.g. differential thresholding without MST can lead to unconnected nodes and as a result networks of different sizes). Afterwards, we investigated network properties over a range of densities (0.036, 0.079, 0.106, 0.132, 0.159, 0.212, 0.238, 0.265, 0.291; percent of all possible connections, i.e. ten thresholding levels) by stepwise adding the highest remaining edges. This lead to ten adjacency matrices, for each frequency band and participant.

To estimate the differences in global network structure of the brain, we analyzed two graph theoretical network parameters reflecting segregation (Average Clustering Coefficient) and integration (Average Shortest Path Length) of the brain [62, 63, 95, 96]. It has been shown in a variety of simulations and network analyses of imaging data, that the human brain, among other biological systems and animal brains [93, 97], exhibits a small world architecture [93], which leads to an advantage of efficient information transfer while keeping the anatomical costs low [98, 99]. Compared to the two studies by Jäncke et al. [41] and Loui et al. [43] the present investigation did use network measures averaged over the whole brain and compared to those of a random network, instead of individual values per region. This is advantageous, as the vast variability of individual coherence within a network is reduced to one value per parameter and participant that reflects the small-worldness or efficiency of a brain network relative to a random or chaotic network [62, 82, 85, 92–94]. By definition [62, 97, 98] Small-Worldness σ (7) is characterized by a C, which is much higher than that of a random network (γ = C real /C random >>1), but has a comparable short path length (λ = L real/L random ≈ 1).
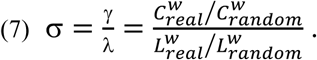

Here the (8) Clustering Coefficient *C_i_* of a node i is defined as the weighted average amount of (9) triangles 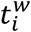 around it, i.e. the sum of connections between the neighbours of a node i, divided by the total amount of possible connections among its neighbors:
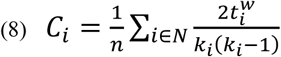

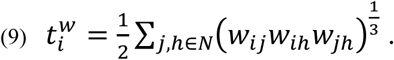

The (10) Global Clustering Coefficient of a weighted association matrix C^w^ denotes the average clustering coefficient summed over all nodes *i* ∊ *N* in a network
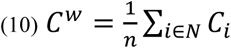

and is interpreted as a measure of segregation of the network.

On the other hand the (11) Characteristic Path Length *L_i_* of a node i is defined as the (12) average pairwise distance 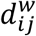 between the node i and any other node j in the weighted (w) network
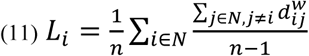

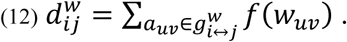

The (13) Global Average Path Length is then calculated by taking the average of the Characteristic Path Length of all nodes *i* ∊ ***N*** in the network
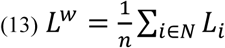

and is interpreted as a measure of integration of the network. As both, γ and λ reflect the underlying brain network structure relative to a random network of the same density (and degree distribution) and influence the calculation of small-worldness, we chose to look at these parameters separately. That is, because we were specifically interested in the potentially differential relation of segregation and integration in the brain. Various authors have shown, that long-range-connections (integration) are more associated with synchronization in low frequency bands, whereas short-range-connectivity is mainly processed within beta-band (e.g. [100]).

### Statistical Analysis

All statistical analyses were done using the open-source statistical software package R (Version 3.5,https://www.r-project.org/).

We expected group differences between AP and RP regarding AQ-Scores, MAD (PAT), PIS (sum of correctly identified tones) and network parameters γ and λ (in beta, delta and theta band). Additional unexpected results obtained in other frequency bands and network parameters are also reported. In order to correct for multiple comparisons across frequency bands, ten thresholds each and various network parameters, only significant results within at least two successive thresholds were considered significant. Results were obtained using t-tests and non-parametric equivalents when applicable. Inter-correlations between the variables were investigated to further explore the interrelation of autistic traits, absolute pitch performance and network structure using regression and bivariate correlations. Finally, the network parameters λ and γ, AQ-Score and the age of beginning to play a musical instrument (as a covariate) were used to predict PIS and PAT performance within the sample using multiple regressions and AQ and AP performance to predict network parameters.

## Results

### Behavioral performance and autism traits

Welch two-sample t-tests revealed significant lower absolute deviations from target tone (MAD; t(43.7)= 15.614; *p* < 2.2e-16) and lower deviations from individual mean deviation, i.e. interpreted as pitch template (SDfoM; t(40.9)= 12.145; *p* = 3.788e-15) for absolute pitch compared to relative pitch possessors (Table 2). Having AP was further associated with more autistic traits (AQ; t(60.3) = −2.501; p < 0.015) and (marginally) an early start of musical training (starting age; t (55.4) = 1.751; p < 0.086). For AQ, only the subscale “imagination” reached significance (t(57.4)=-4.287, p < 6.997e-05) with higher values for AP, while “communication” (t(55.3)=-1.977, p = 0.053) and “attention to detail” (t(61.6)=-1.776, p = 0.081) were marginally and “social skills” (t(60.9)=- 1.145, p = 0.257) and “attention switching” (t(62.0)=1.012, p = 0.316) not significant.

**Table 2.** Group differences

|  | AP (n=31) |  |  | RP (n=33) |  |  | t-test* |
| --- | --- | --- | --- | --- | --- | --- | --- |
|  | Mean | SD | Range | Mean | SD | Range |  |
| **AQ** | 20.48 | 6.05 | 10-36 | 16.88 | 5.44 | 6-27 | **t(60.3) = -2.501;**<br>**p = 0.015** |
| **MAD** | 41.37 | 36.49 | 9.8 -<br>200.57 | 296.84 | 86.12 | 91.04 -<br>467.52 | **t(43.7) = 15.614;**<br>**p < 2.2e-16** |
| **SDfoM** | 52.31 | 44.96 | 7.41-235.69 | 329.77 | 122.77 | 134.37 -<br>811.73 | **t(40.9) = 12.145;**<br>**p = 3.788e-15** |
| starting age | 5.97 | 2.97 | 2-17 | 7.12 | 2.22 | 3-12 | t(55.4) = 1.751;<br>p = 0.086 |

Age, nonverbal IQ (SPM), information processing capacity (ZVT), musical training (total hours during life span on main instrument), musicality (AMMA; MSI) and online pitch identification screening (PIS) for each group; \* one RP has reported himself not having absolute pitch but reached a screening score of 13. Because of this and the weak performance in the pitch adjustment test, the subject was assigned to the RP group. Significant group differences highlighted in bold. \* Welch-two sample t-test

### Network analysis

Welch two sample t-tests (p<0.05, uncorrected) reveiled higher average Path length λ for AP compared to RP within the delta band (2-4 Hz) for both, eyes open (EO) and eyes closed (EC), resting state conditions and at least two thresholds each. Lower path length values for AP were found in alpha (7-13 Hz) and beta (13-18 Hz) eyes open condition for one threshold each but did not reach significance (p<0.10; see figure 1). Analysis of Clustering Coefficient γ yielded lower Clustering for AP in EO delta (p<0.05) for one threshold and EO beta (p<0.10) for two neighboring thresholds. RP exhibited higher Clustering for a single threshold in EO theta (p<0.10). Small Worldness σ was widely reduced in AP within EC gamma, EC alpha and EO alpha with significant (p<0.05) or marginal significant (p<0.10) group differences across one or two thresholds each (figure 1). No significant higher thresholds were found for AP.

**Figure 1:**
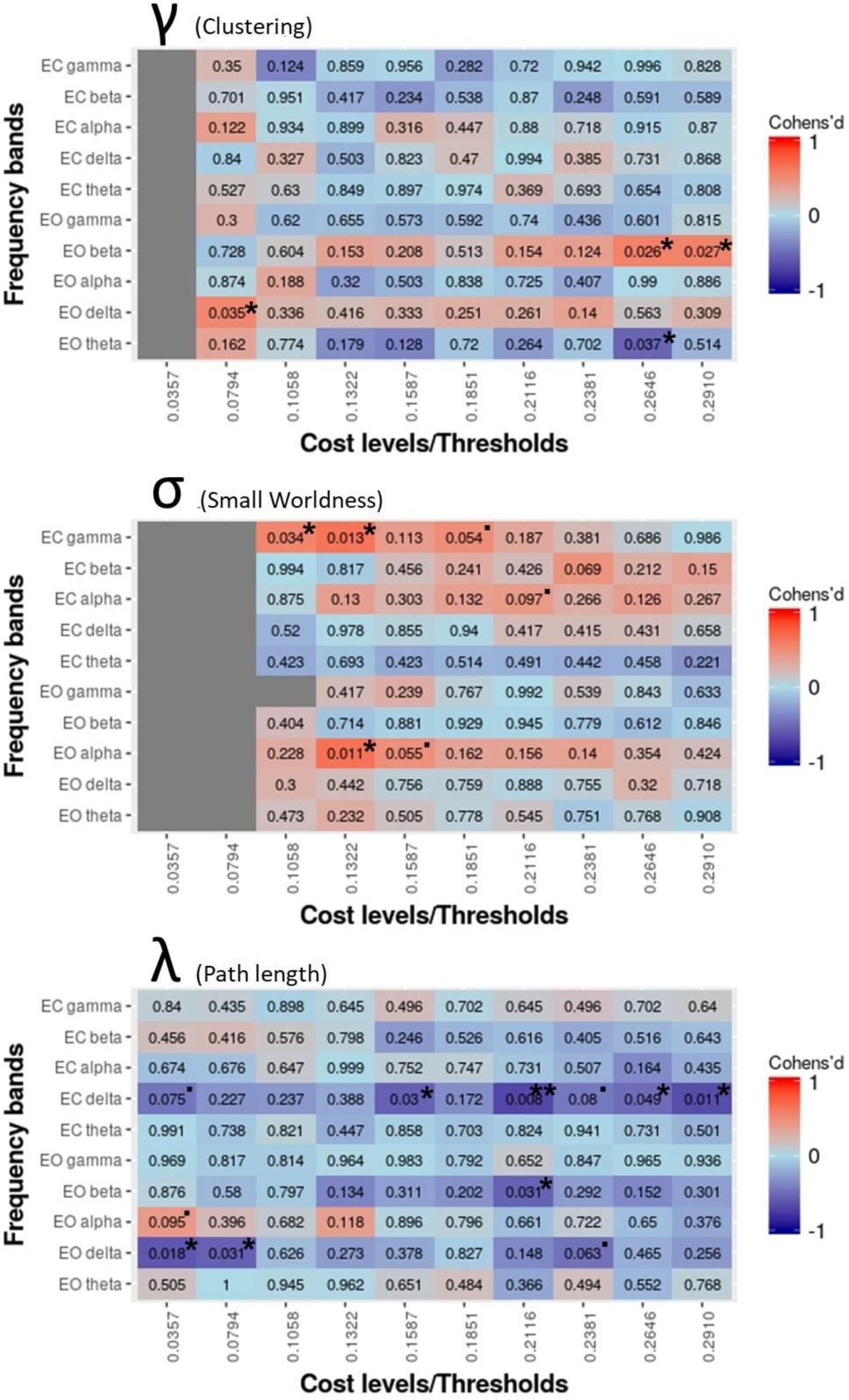
Multiple comparisons (Welch two sample t-tests) across frequency bands, thresholds, eyes-closed vs. eyes-open RS between AP and RP. Matrix cells contain p-values (uncorrected) and are colored according to Cohen’ d values. Blue cells indicate higher SW (Small World), Lrand (Path length compared to random network) and Crand (Clustering compared to random network) for AP compared to RP; red cells show higher parameters for RP. Significant results (p<0.05, *; p<0.01,**; p<0.001,***) and tendencies (p<0.10, “.”) are marked.

In general, significant and marginally significant results were spread widely across different thresholds (see figure 1). Only significant results appearing on at least two thresholds in the same frequency band were included in further analyses (multiple regression). Of those, the threshold (T) with the highest effect size of neighbouring significant results was taken: **Clustering γ EO beta (T= 0.2910), Small Worldness σ EC gamma (T=0.1322) and Path length λ EC delta (T=0.2116).** Path length EO delta (T=0.0357) was not taken into account because of correlation with Path length EC delta (T=0.2116).

### Prediction of absolute pitch performance

Multiple regression analysis was used to predict AP performance in pitch naming (PIS) and pitch adjustment (PAT). A multiple regression predicting PIS performance by autistic traits (AQ; beta = 0.892, p<0.0001), Clustering (C_EO_b10; beta = −66.074, p<0.0002), Path length (L_ECd10; beta = 76.909, p<0.008) and Small Worldness (ECg4; beta= −6.612,p<0.0325) explained 44% of the variance (R^2^ = 0.44, R^2^_adjusted_=0.401; F(4,57) = 11.22, p <9.92e-17). PAT performance was predicted by the same predictors plus the age of begin of musical training (starting age) and explained 38% of the variance (R^2^ = 0.380, R^2^_adjusted_=0.326; F(5,57) = 6.991, p <3.736e-05). Here, AQ (beta = −0.089, p<0.004), Clustering (beta = 6.775, p<0.004) and Small Worldness (beta= 0.946, p<0.023)) significantly contributed to the prediction, while age of begin of musical training (beta= 0.130, p<0.053)) and Path length (beta= −7.006, p<0.070)) remained marginal significant. Bivariate pearson correlations among the variables are listed in table 3.

**Table 3.** Bivariate correlations between variables of interest

| Correlation coefficient (Pearson) |  |  |  |  |  |  |  |  |
| --- | --- | --- | --- | --- | --- | --- | --- | --- |
|  | **PIS** | 0.38** | -0.91*** | -0.85*** | -0.23 . | 0.35** | -0.30* | -0.28* |
|  | 0.002** | **AQ** | -0.28* | -0.25* | 0.025 | 0.13 | 0.20 | 0.022 |
|  | <0.001*** | 0.024* | **MAD<sup>a</sup>** | 0.93*** | 0.30* | -0.27* | 0.28* | 0.31* |
| p-value | <0.001*** | 0.045* | p<0.001*** | **SDfoM<sup>a</sup>** | 0.25 . | -0.21 . | 0.25 . | 0.21 . |
|  | 0.074 . | 0.844 | 0.017* | 0.053 . | **start age** | 0.018 | 0.22 . | 0.10 |
|  | 0.005** | 0.315 | 0.033* | 0.094 . | 0.887 | **$\lambda$ ECdelta** | 0.08 | -0.23 . |
|  | 0.017* | 0.109 . | 0.026* | 0.051 . | 0.079 . | 0.534 | **$\gamma$ EObeta** | -0.044 |
|  | 0.028* | 0.866 . | 0.013* | 0.100 . | 0.431 | 0.075 . | 0.731 | **$\sigma$ ECgamma** |

Pearson correlations between variables of interest (network parameters: selected bands and thresholds); significant correlation coefficients are highlighted with stars. <sup>a</sup> variables were z-standardized to the mean and sd of the non-AP population

### Prediction of network parameters

To further investigate the interrelation between AP, autistic traits and network connectivity, we calculated general linear models to predict network connectivity (L, C, SW) differences obtained before by a combination of AP performance and AQ. Different models were compared using R^2^, R^2^_adjusted_ and information criteria (AIC). Separate models are shown for active (PAT) and passive (PIS) AP performance as for their high collinearity. Only Clustering obtained a better prediction by a joint model of AQ and AP performance (active and passive on separate models because of intercorrelation) with AQ as a significant predictor. While inclusion of AQ-Scores did not improve the prediction of path length and small worldness (see table 4), it was predictive for Clustering Coefficients in the beta range in each, a joint model with either MAD (F(2,60)=6.011, p<0.004; R^2^=0.167, R^2^_adjusted_; β_AQ_=4.06e-3, p<0.014; β_MAD_=2.07e-4, p<0.004) or PIS-performance (F(2,59)=6.889, p<0.002; R^2^=0.189, R^2^_adjusted_; β_AQ_=4.44e-3, p<0.009; β_MAD_=-2.62e-3, p<0.0041). Both models were superior compared to a prediction of network connectivity by AP performance alone, even though the bivariate correlation between AQ and Clustering did not reach significance (see previous section)

**Table 4.**
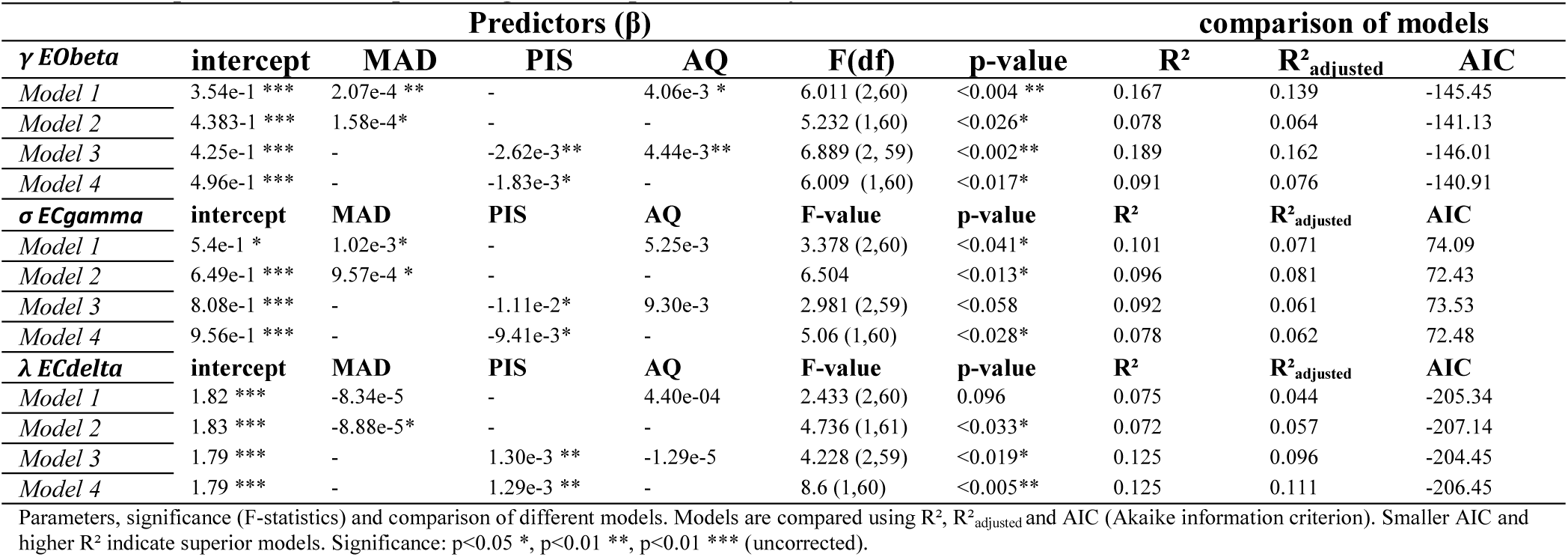
Comparison of models predicting network parameters by AP and AQ

| $\gamma$ EObeta | Predictors ($\beta$) | | | | comparison of models | | | | |
| --- | --- | --- | --- | --- | --- | --- | --- | --- | --- |
|  | intercept | MAD | PIS | AQ | F(df) | p-value | R<sup>2</sup> | R<sup>2</sup><sub>adjusted</sub> | AIC |
| Model 1 | 3.54e-1 *** | 2.07e-4 ** | - | 4.06e-3 * | 6.011 (2,60) | <0.004 ** | 0.167 | 0.139 | -145.45 |
| Model 2 | 4.383e-1 *** | 1.58e-4 * | - | - | 5.232 (1,60) | <0.026 * | 0.078 | 0.064 | -141.13 |
| Model 3 | 4.25e-1 *** | - | -2.62e-3 ** | 4.44e-3 ** | 6.889 (2, 59) | <0.002 ** | 0.189 | 0.162 | -146.01 |
| Model 4 | 4.96e-1 *** | - | -1.83e-3 * | - | 6.009 (1,60) | <0.017 * | 0.091 | 0.076 | -140.91 |
| $\sigma$ ECgamma | intercept | MAD | PIS | AQ | F-value | p-value | R<sup>2</sup> | R<sup>2</sup><sub>adjusted</sub> | AIC |
| Model 1 | 5.4e-1 * | 1.02e-3 * | - | 5.25e-3 | 3.378 (2,60) | <0.041 * | 0.101 | 0.071 | 74.09 |
| Model 2 | 6.49e-1 *** | 9.57e-4 * | - | - | 6.504 | <0.013 * | 0.096 | 0.081 | 72.43 |
| Model 3 | 8.08e-1 *** | - | -1.11e-2 * | 9.30e-3 | 2.981 (2,59) | <0.058 | 0.092 | 0.061 | 73.53 |
| Model 4 | 9.56e-1 *** | - | -9.41e-3 * | - | 5.06 (1,60) | <0.028 * | 0.078 | 0.062 | 72.48 |
| $\lambda$ ECdelta | intercept | MAD | PIS | AQ | F-value | p-value | R<sup>2</sup> | R<sup>2</sup><sub>adjusted</sub> | AIC |
| Model 1 | 1.82 *** | -8.34e-5 | - | 4.40e-04 | 2.433 (2,60) | 0.096 | 0.075 | 0.044 | -205.34 |
| Model 2 | 1.83 *** | -8.88e-5 * | - | - | 4.736 (1,61) | <0.033 * | 0.072 | 0.057 | -207.14 |
| Model 3 | 1.79 *** | - | 1.30e-3 ** | -1.29e-5 | 4.228 (2,59) | <0.019 * | 0.125 | 0.096 | -204.45 |
| Model 4 | 1.79 *** | - | 1.29e-3 ** | - | 8.6 (1,60) | <0.005 ** | 0.125 | 0.111 | -206.45 |

Parameters, significance (F-statistics) and comparison of different models. Models are compared using R<sup>2</sup>, R<sup>2</sup><sub>adjusted</sub> and AIC (Akaike information criterion). Smaller AIC and higher R<sup>2</sup> indicate superior models. Significance: $p<0.05$ \*, $p<0.01$ \*\*, $p<0.01$ \*\*\* (uncorrected).

### Post-hoc analysis: single connection statistics

To assess single connection differences in the beta frequency band, permutation statistics (n_permutations_=10000) across groups were evaluated post-hoc. To obtain these, raw matrices in the relevant frequency bands (significant results) were z-standardized individually and permutation group statistics (FDR corrected) performed across groups using custom MATLAB scripts. An unstandardized comparison was provided as well. While the former reflects the relative importance of the connections within the participants’ networks between the groups, the latter shows group differences in the absolute wPLI. Results revealed overall increased wPLI values for AP in a network comprising mainly left frontal and parietal regions (especially nodes: F7, F3, F4, P3; see table 5 for anatomical correlations) combined with lower connectivity within and between bilateral temporal regions (FT7-T8, FT7-T7, FT8-T8; unstandardized results). Relative to their own networks (z-standardized participants matrices), AP’s exhibited reduced connectivity compared to RP between left FT7 and various sites along frontal-temporal-occipital electrodes (F8 ,T8,TP8,P8,P3) in the right hemisphere, especially again within and between bilateral temporal regions (FT7-T8, FT7-T7, FT8-T8). The only significant higher connections relative to their own network for AP were found between F7, F8 and P7. Figure 2 (brain nets created using the Matlab Toolbox BrainNet Viewer [101]) shows Cohen’s d effect size values for all pairs of electrodes between groups in separate matrices for z-standardized vs. unstandardized raw connectivity matrices. The most pronounced differences that were found in both, standardized and unstandardized (relative) comparisons, comprise reduced interconnection between bilateral auditory cortices (FT7-T8, FT7-T7, FT8-T8) as well as higher frontal-parietal connectivity (F7-F8, F8-P7) for AP. These connections therefore not only exhibit a group difference on absolute wPLI values, but also play a different role relative to the other connections in the participants networks.

**Table 5.** Cranio-Cerebral Correlations for electrode positions (10-10 system, modified after [110])

| Electrode label | Talairach coordinates (mm) |  |  | lobe | gyri | anatomical region |
| --- | --- | --- | --- | --- | --- | --- |
|  | x | y | z |  |  | BA |
| FP1 | -21.2±4.7 | 66.9±3.8 | 12.1±6.6 | L FL | Superior frontal G | 10 (100%) |
| FP2 | 24.3±3.2 | 66.3±3.5 | 12.5±6.1 | R FL | Superior frontal G | 10 (100%) |
| F3 | -39.7±5.0 | 25.3±7.5 | 44.7±7.9 | L FL | Middle frontal G | (75%), 6 (19%), 46 (6%) |
| F4 | 41.9±4.8 | 27.5±7.3 | 43.9±7.6 | R FL | Middle frontal G | 8 (69%), 6 (6%), 9 (25%) |
| FC3 | -45.5±5.5 | 2.4±8.3 | 51.3±6.2 | L FL | Middle frontal G | 6 (75%), 4 (12,5%), 8 (12,5%) |
| FC4 | 47.5±4.4 | 4.6±7.6 | 49.7±6.7 | R FL | Middle frontal G | 8 (69%), 6 (6%), 9 (25%) |
| C3 | -49.1±5.5 | -20.7±9.1 | 53.2±6.1 | L PL | Postcentral G | 21 (62,5%), 22 (25%), 20 (6,5), 42 (6%) |
| C4 | 50.3±4.6 | -18.8±8.3 | 53.0±6.4 | R PL | Postcentral G | 123 (81,5%), 6 (12,5), 40 (6%) |
| CP3 | -46.9±5.8 | -47.7±9.3 | 49.7±7.7 | L PL | Inferior parietal G | 40 (82%), 123 (6%), 5 (6%), 39 (6%) |
| CP4 | 49.5±5.9 | -45.5±7.9 | 50.7±7.1 | R PL | Inferior parietal G | 40 (77,5%), 123 (12,5%) |
| P3 | -41.4±5.7 | -67.8±8.4 | 42.4±9.5 | L PL | Precuneus | 39 (37,5%), 7 (25%), 19 (25%), 40 (12,5%) |
| P4 | 44.2±6.5 | -65.8±8.1 | 42.7±8.5 | R PL | Inferior parietal L | 39 (31%), 7 (25%), 40 (25%), 19 (19%) |
| F7 | -52.1±3.0 | 28.6±6.4 | 3.8±5.6 | L FL | Inferior frontal G | 45 (56%), 47 (38%), 46 (6%) |
| F8 | 53.2±2.8 | 28.4±6.3 | 3.1±6.9 | R FL | Inferior frontal G | 45 (37,5%), 47 (37,5%), 46 (25%) |
| FT7 | -59.2±3.1 | 3.4±5.6 | -2.1±7.5 | L TL | Superior temporal G | 22 (75,5%), 21 (12,5%), 38 (6%), 44 (6%) |
| FT8 | 60.2±2.5 | 4.7±5.1 | -2.8±6.3 | R TL | Superior temporal G | 22 (75%), 21 (13%), 38 (6%), 44 (6%) |
| T7 | -65.8±3.3 | -17.8±6.8 | -2.9±6.1 | L TL | Middle temporal G | 21 (81,5%), 22 (12,5%), 43 (6%) |
| T8 | 67.4±2.3 | -18.5±6.9 | -3.4±7.0 | R TL | Middle temporal G | 4 (50%), 123 (25%), 6 (25%) |
| TP7 | -63.6±4.5 | -44.7±7.2 | -4.0±6.6 | L TL | Middle temporal G | 21 (50%), 37 (25%), 22 (19%), 20 (6%) |
| TP8 | 64.6±3.3 | -45.4±6.6 | -3.7±7.3 | R TL | Middle temporal G | 21 (62,5%), 22 (12,5%), 20 (12,5%), 37 (12,5%) |
| P7 | -55.9±4.5 | -64.8±5.3 | 0.0±9.3 | L TL | Inferior temporal G | 37 (44%), 19 (38%), 39 (18%) |
| P8 | 56.4±3.7 | -64.4±5.6 | 0.1±8.5 | R TL | Inferior temporal G | 19 (56%), 37 (19%), 20 (12,5), 39 (12,5%) |
| O1 | -25.8±6.3 | -93.3±4.6 | 7.7±12.3 | L OL | Middle occipital G | 18 (81%), 19 (19%) |
| O2 | 25.0±5.7 | -95.2±5.8 | 6.2±11.4 | R OL | Middle occipital G | 18 18 (81%), 19 (19%) |
| Oz | 0.3±5.9 | -97.1±5.2 | 8.7±11.6 | M OL | Cuneus | 18 (62,5), 17 (31%), 19 (6,5%) |
| Fz | 0.0±6.4 | 26.8±7.9 | 60.6±6.5 | M FL | Bilateral medial | 6 (81,5%), 8 (12,5%), 9 (6%) |
| Cz | 0.8±4.9 | -21.9±9.4 | 77.4±6.7 | M FL | Precentral G | 4 (62,5%), 6 (37,5%) |
| Pz | 0.7±6.3 | -69.3±8.4 | 56.9±9.9 | M PL | Superior parietal L | 7 (88%), 5 (6%), 19 (6%) |

Estimated projection of electrode positions to cortical areas (Talairach space) and variability of associated BA (Brodman areas), investigated by [110] using EEG-MRI sensors. L=left, R=right, FL= frontal lobe, PL=parietal lobe, TL=temporal lobe, OL=occipital lobe; L=lobe, G=Gyrus.

**Figure 2.**
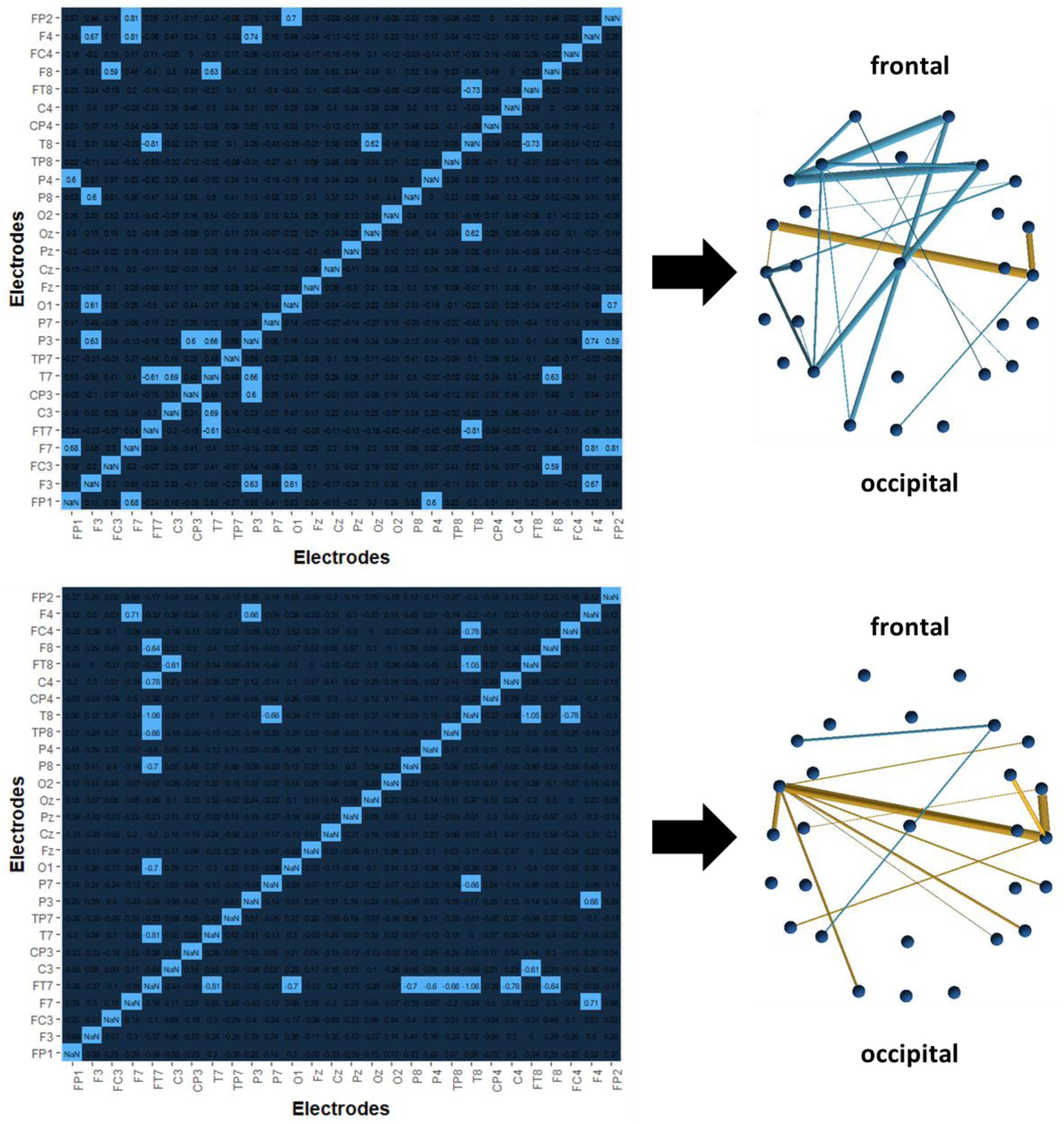
Visualization of single connection differences in the beta range. left: Cohen’s d effect size values for all pairs of electrodes between groups in separate matrices for unstandardized (top) vs. z-standardized (bottom) raw connectivity matrices (permutation testing). Significant connections (FDR corrected) are highlighted in light blue. right: significant differences plotted in EEG-cap order (extended 10-20 system, view from above). Colors indicate direction of effect (blue: AP>RP, yellow: RP<AP) and size of the line the corresponding effect size (Cohen’s d).

Rough anatomical associations of electrode positions, taken from Koessler et al. [102], are summarized in table 5. However, it must be clearly said, that graph theoretical accounts and single connection permutation tests are completely different techniques and cannot be compared directly. This is, because in the course of graph theoretical analysis, thresholds have to be applied on the participants’ raw matrices, leading to a reduced number of total connections. Thus the connections fed into graph analysis also highly depend on the participant specific order of connection weights and can have a high regional variability despite producing similarly high or low network parameters.

## Discussion

The results of the present study underline a possible interrelation between autistic traits, brain connectivity and absolute pitch ability. We investigated EEG resting state connectivity using a graph theory approach in professional musicians with and without absolute pitch, the Autism Spectrum Quotient [70] and each a test of pitch naming and pitch adjustment ability. Analyses revealed higher autistic traits, higher average Path length (delta 2-4 Hz)), lower average Clustering (beta 13-20 Hz), lower Small-Worldness (gamma 30-60 Hz) and a tendency for an earlier start of musical training in absolute pitch musicians. Furthermore, pitch naming was well predicted by autistic traits, Path length and Clustering values, explaining a total of 44% of the variance. Pitch adjustment (i.e. active absolute pitch) was explained by the same predictors plus the age of begin of musical training summing up to an R^2^ = 0.38. However, in the latter case, the starting age of musical training and Path length remained marginally significant.

It is noteworthy that the start of playing a musical instrument in our models did not significantly improve the prediction of AP performance but only in pitch adjustment. Furthermore, the total amount of musical training during life was neither predictive of any AP performance in the general linear model, nor did show a group difference. The typical human brain exhibits a small-world like structure with a much higher Clustering compared to a random network, while maintaining an efficient information transfer and low wiring cost through an equally low path length [62, 93, 97]. In this context, the results of the present study indicate a less efficient, less small-world structured functional network in AP compared to RP, in line with the structural results of Jäncke et al. [41] and results from the autism research [44, 45, 48, 90, 103] but extends the results to EEG functional connectivity networks.

It is further interesting that both correlations and regressions between autistic traits and the two AP test show higher correlations and better prediction of pitch naming than pitch adjustment by AQ. This can be explained by the aforementioned theory of veridical mapping [7, 61]. This framework explains savant abilities and other unusual abilities in autism by their common characteristic of one-to-one mappings between elements of two conceptual or perceptual structures (e.g. letters-musical tones, letters-colors). According to this theory all of these abilities share further commonalities including hyper-systemizing [53], enhanced perceptual functioning [51, 52], depend on exposure to material and - if they occur as autistic savant ability - the related elements can also be recalled without a strategy [7, 61]). This explicit recall in absolute pitch, i.e. the naming of the pitch, therefore might be a more savant-like ability, leading to a higher correlation with autistic traits.

Furthermore, we observed reduced connectivity for AP compared to RP in interhemispheric connections when compared to the participants own distribution of connectivities (z-standardized calculation) – especially between left auditory located electrodes and various right temporal, parietal and frontal electrodes.

While higher Path length in low frequency bands (delta, therefore reduced integration) and lower Clustering in higher frequencies (beta, reduced segregation on sensor level) are in line with our apriori hypotheses, we did not expect reduced Small-Worldness within gamma-band for AP compared to RP (found during eyes closed). Nevertheless this result can be explained by previous research findings: Cantero et al. [104] reported increased gamma band measured by intracranial electrodes between hippocampal areas and neocortex in humans during wakefulness but not during sleep, pointing to a relation of gamma-band couplings and awareness states in humans. This also suggests, that gamma band activity, probably useful for the storage and retrieval of memory [105–107] and binding of perceptual features [106, 107] might even play a role during resting (awake more than asleep) states. AP ability, similarly, is often described as the ability to associate tones and verbal labels in a stable, hyper memorized way, pointing to the importance of long-term memory processes [108–112]. Furthermore, Bhattacharya et al. [113, 114] found increased long-range gamma synchronization between distributed cortical areas during music listening in musicians compared to non-musicians, which might reflect musical memory and binding of musical features. In contrast, Sun et al. [115] found reductions in gamma band phase locking and power in participants with autism associated with perceptual organization tasks (visual), while Brown et al. [116] found higher gamma peaks in response to illusory figures in autism. Generally, abnormal gamma activity is found in a range of neuropsychiatric disorders, with reduced gamma in negative schizophrenic symptoms, Alzheimer’s disease and task specific gamma decrease in autism, but an increase in gamma in ADHD, positive schizophrenic symptoms and epilepsy (for a review see [117, 118]). Thus, the results of reduced Small-Worldness in AP are in line with an integration-deficit hypothesis of AP, both in perceptual organization and binding of musical stimuli and in brain connectivity, which is again similar to autism (see [42, 44, 119–122]. However, the findings in gamma band did not show correlations with autistic symptoms.

Our results replicate the results of Dohn et al. [39] showing higher autistic traits, which reached significant in the subscales “imagination” (similar to [39]),“attention to detail” (marginally) and “social skills” (marginally). Furthermore, autistic traits were also not only correlated to pitch naming as already shown by Dohn et al. [39], but also to pitch adjustment accuracy (MAD, mean absolute deviation to target tone in cent ; 100 cent= 1 semitone) and adjustment consistency (SDfoM, pitch template tuning). However, similar to [39], group mean autistic traits did not reach the cutoff for diagnostic relevance, indicating a high variability regarding autistic traits even in the AP group (with 7 AP compared to 1 RP scoring above cutoff or borderline). This fits with analyses of the broader autism phenotype [123] and might implicate joint as well as divergent phenotypic and endotypic characteristics of AP and autism.

In contrast to our study, various previous studies have shown an influence of the start of musical training in AP, making the onset of training before the age of 7 necessary, but not sufficient to acquire absolute pitch [12, 16–19, 36]. For example, Loui et. al [36] recently found, that early onset of musical training was associated with an enlarged tract between pSTG and pMTG in the left hemisphere, but the degree of AP proficiency still correlated with the size of the tract after partialling out age of onset. Gregersen et al. [12] further analyzed familiar aggregation of AP in different samples of musicians and non-musicians with early and late onset of musical training comparing different types of musical education and found no general differences of AP between early or late starting siblings of AP. Their results further indicated a higher influence of genetic disposition and the type of education used, which both had a more pronounced influence than age of onset per se [12].

Higher average Path length (delta 2-4 Hz)), lower average Clustering (beta 13-20 Hz) and lower Small-Worldness (gamma 30-60 Hz) for AP compared to RP are also in line with previous studies showing structural local hyper-vs. global hypoconnectivity in AP [41] and reduced Clustering and higher Path length in participants with autism [103, 134]. In contrast, Loui et al. [43] reported overall increased degrees, clustering and local efficiency coefficients of functional networks in AP using fMRI during music listening and rest. The authors further speculate that there might be a “dichotomy between structural and functional hyperconnectivity in AP, where structure is locally hyperconnected but function is globally hyperconnected [43]. The present study, however, provided more evidence for an also functionally underconnected brain in AP musicians compared to relative pitch musicians. Diverging results compared to Loui et al. [43] might be due to differences in methods (EEG vs. fMRI) or different definition of nodes (electrode positions vs. brain regions) and edges (wPLI vs. functional correlations).

Differences seen in single connection analysis might reflect the connections that lead to differences in Clustering values described above. Similarly to the prediction of Clustering by AP and autistic traits, single connection differences in the beta range are in line with findings from the autism literature: First, various others have reported reduced interhemispheric connectivity in autism [48, 102, 103, 135, 136]. Second, hypoconnectivity between left FT7 (BA: 22) and right frontal-temporal-occipital electrodes (F8, T8, TP8, P8, P4; BA: 45/47, 4, 21/22/20/37, 19/37, 39/7/40/19; see table 4 for anatomical interpretation of electrode positions) might reflect a specific underconnectivity between left STG and right IFOF, of which alterations have already been described in both AP [137] and Autism [138]. Especially reduced interhemispheric connectivity between left auditory related cortex and right IFOF might reflect autism-like personality traits and perception of (some) absolute pitch possessors. The IFOF, especially the right IFOF, has been shown to play an important role in music perception and the integration of musical features, as it connects various brain regions from frontal over temporal to posterior parts of the brain [139]. A reduced white matter integrity of IFOF was found in amusics [139, 140], whereas people with synaesthesia and absolute pitch where shown to have a higher IFOF integrity [59, 137]. More importantly, however, increased interhemispheric connectivity in musicians was found by several studies [141–145] showing the importance of interhemispheric integration in music perception. A reduced interhemispheric functional connectivity, especially between bilateral auditory regions as found in the present study, perhaps might result in less perceptual integration of musical features (i.e. auditory weak central coherence) and hence a more detail oriented processing of music and musical pitches (i.e. absolute vs. relative) in those participants. An exaggeration of those features might also lead to symptoms of amusia, which has also been associated with alterations in left and right STG and right IFOF [139, 140, 146] and with autism [147]. However, it must be clearly said, that we cannot explicitly conclude anatomical differences from connectivity differences on the sensor levels. Further structural or functional studies using methods with high anatomical precision have to be conducted to evaluate this hypothesis.

Some caveats of the present approach are warranted. First, we did not use a source-based approach of functional connectivity, making conclusions with respect to anatomical associations of the obtained differences very speculative. Second, various different configurations of local and global hyper- vs. hypoconnectivity can be assumed to result into the same averaged network measures, therefore no conclusions can be made about the exact relative structure within the brain and among different regions. Nevertheless, higher Path length (EC, delta 2-4 Hz) can be interpreted as weaker integration in the network and higher Clustering (EO,13-20 Hz) as higher local segregation of functions [85] and therefore might again reflect a local hyper- over global (integrative) hypoconnectivity in the brain of AP musicians. This interpretation is further encouraged by studies showing, that long-range connectivity (integration) is more reflected in low frequency bands, whereas short range connectivity is more high frequency bands [100, 148]. This again fits to the results of our study, as higher Clustering, indicative for local segregation, was found in the beta range and Path length - indicative for global integration in the network and therefore long-range-associations - in the delta range.

In addition, significant group differences were highly selective for certain frequency bands, states (EO vs. EC) and thresholds. Nevertheless, we can rule out the possibility, that we obtained those differences by chance. First, there were significant differences for at least one threshold in a frequency band, effect sizes of the other thresholds in the same frequency band never (exclusive: Crand EO alpha) indicated reverse effects (see color code in figure 1). Second, we did only consider differences relevant, if at least two neighbouring thresholds exhibited a significant group difference. Third, the three network parameters selected via group differences always could also predict AP performance with a reasonable high R^2^ and/or showed bivariate correlations with AP performance in both tests of AP.

For the first time we included a pitch adjustment test of active absolute pitch [69] into a study on brain connectivity in AP, so we are not only referring to pitch naming as were previous studies [36, 39, 41, 43]. Also, whereas Jäncke et al. [41] were using structural cortical thickness covariations and Loui et al. [43] functional correlations of fMRI activity (during rest and music listening) as weights for connections in graph analysis, we for the first time applied graph theory on resting state EEG connectivity of AP musicians, both in eyes closed and eyes open conditions. This is similar to methods used in analyzing brain connectivity in autism [49, 103]. Finally, while e.g. Elmer et al. [108] used phase synchronization as an estimate for functional EEG connectivity, we used wPLI (weighted phase lag index, [75]), which is less contaminated by volume conduction [75–78, 81] thus contributing to a higher validity and reliability with respect to true brain connectivity and graph theoretical parameters [79, 83, 84].

In summary, differences in network and connectivity analysis in the beta band seem to be specifically associated with the relation of autistic traits and absolute pitch, whereas Path length in delta range and Small-Worldness in gamma range might reflect other influences on the acquisition of the ability (e.g. environmental factors, genetic factors not attributable to autistic traits, musical education method, instrument, learning, sensitive periods). To our knowledge this is the first study to combine measures on autistic traits and brain networks on musicians with and without absolute pitch. We conclude that this is further evidence showing, that AP and Autism both have shared and distinct neuronal and phenotypic characteristics. This might also be reflected in subgroups of AP with different genesis, providing new arguments for the discussion about a dichotomous or continuous view on AP. However, the causal relationship between AP, autistic traits and brain connectivity remains to be evaluated.

## List of abbreviations

EEG: electroencephalography
(w)PLI: (weighted) Phase Lag Index
AP: absolute pitch
RP: relative pitch
ASD: Autism Spectrum Disorder or Condition
PIS: Pitch identification Screening
SPM: Standard Progressive Matrices
ZVT: “Zahlenverbindungstest” (~Trail Making Test)
AMMA: Advanced Measures of Music Audiation
(GOLD-)MSI: Musical-Sophistication Index
PAT: Pitch adjustment test
MAD: Mean absolute derivation from target tone
SDfoM: Standard deviation from own mean
AQ: Autism-Quotient
EO: eyes open resting state
EC: eyes closed resting state
ICA: independent component analysis
PC: phase coherence
sgn: sign
imag: imaginary component
S_xyt_: cross-spectrum
MST: minimum spanning tree
σ: Small-Worldness
C: Clustering Coefficient
γ: Clustering relative to random network of same cost and density distribution
L: Path length
λ: Path length relative to random network of same cost and density distribution

## Declarations

**Ethics approval and consent to participate:** the study was approved by the ethic committee of the Hanover Medical School (Approval no. 7372, committee’s reference number: DE 9515). All participants gave written consent.

**Consent for publication:** not applicable

### Availability of data and material

The datasets generated and/ or analysed during the current study are not publicly available due to specifications on data availablity within ethics approval. Data are however available from the corresponding author upon reasonable request and with permission of the ethics committee of the Hanover Medical School.

### Authors’contributions

TW designed the study, collected, analysed and interpreted the data. RB made intensive contributions to preprocessing of EEG data, analysis of network parameters and single connection differences as well as provided further ideas on data analysis and interpretation. SBC contributed to the interpretation of the results and improvement of the manuscript. EA contributed to the design of the study and interpretation of the data. All authors read, improved and approved the final manuscript.

## Acknowledgements

Hannes Schmidt, Pablo Carra, Artur Ehle, Fynn Lautenschläger, Michael Großbach, Christos Ioannou. SBC and RB were supported by the Autism Research Trust and the MRC during the period of this work.

## References

1. Baron-Cohen S, Wheelwright S, Burtenshaw A, Hobson E. Mathematical Talent is Linked to Autism. Hum Nat. 2007;18:125–31.

2. Mitchell P, Ropar D. Visuo-spatial abilities in autism: A review. Infant Child Dev. 2004;13:185–98.

3. Heaton P, Hermelin B, Pring L. Autism and Pitch Processing: A Precursor for Savant Musical Ability? Music Percept Interdiscip J. 1998;15:291–305.

4. Howlin P, Goode S, Hutton J, Rutter M. Savant skills in autism: psychometric approaches and parental reports. Philos Trans R Soc B Biol Sci. 2009;364:1359–67.

5. Bor D, Billington J, Baron-Cohen S. Savant Memory for Digits in a Case of Synaesthesia and Asperger Syndrome is Related to Hyperactivity in the Lateral Prefrontal Cortex. Neurocase. 2008;13:311–9.

6. Stevens DE, Moffitt TE. Neuropsychological profile of an asperger’s syndrome case with exceptional calculating ability. Clin Neuropsychol. 1988;2:228–38.

7. Bouvet L, Donnadieu S, Valdois S, Caron C, Dawson M, Mottron L. Veridical mapping in savant abilities, absolute pitch, and synesthesia: an autism case study. Front Psychol. 2014;5. doi:10.3389/fpsyg.2014.00106.

8. Mottron Sylvie Belleville Emmanuel L. Atypical Memory Performance in an Autistic Savant. Memory. 1998;6:593–607.

9. O’Connor N, Hermelin B. The memory structure of autistic idiot-savant mnemonists. Br J Psychol. 1989;80:97–111.

10. Lai M-C, Lombardo MV, Chakrabarti B, Baron-Cohen S. Subgrouping the Autism “Spectrum”: Reflections on DSM-5. PLoS Biol. 2013;11:e1001544.

11. Takeuchi AH, Hulse SH. Absolute pitch. Psychol Bull. 1993;113:345–61.

12. Gregersen PK, Kowalsky E, Kohn N, Marvin EW. Early childhood music education and predisposition to absolute pitch: Teasing apart genes and environment. Am J Med Genet. 2001;98:280–2.

13. Gregersen PK, Kowalsky E, Kohn N, Marvin EW. Absolute Pitch: Prevalence, Ethnic Variation, and Estimation of the Genetic Component. Am J Hum Genet. 1999;65:911–3.

14. Deutsch D, Henthorn T, Marvin E, Xu H. Absolute pitch among American and Chinese conservatory students: Prevalence differences, and evidence for a speech-related critical perioda). J Acoust Soc Am. 2006;119:719–22.

15. Profita J, Bidder TG, Optiz JM, Reynolds JF. Perfect pitch. Am J Med Genet. 1988;29:763–71.

16. Zatorre RJ. Absolute pitch: a model for understanding the influence of genes and development on neural and cognitive function. Nat Neurosci. 2003;6:692–5.

17. Baharloo S, Johnston PA, Service SK, Gitschier J, Freimer NB. Absolute Pitch: An Approach for Identification of Genetic and Nongenetic Components. Am J Hum Genet. 1998;62:224–31.

18. Athos EA, Levinson B, Kistler A, Zemansky J, Bostrom A, Freimer N, et al. Dichotomy and perceptual distortions in absolute pitch ability. Proc Natl Acad Sci. 2007;104:14795–800.

19. Deutsch D, Dooley K, Henthorn T, Head B. Absolute pitch among students in an American music conservatory: Association with tone language fluency. J Acoust Soc Am. 2009;125:2398–403.

20. Gregersen PK, Kowalsky E, Lee A, Baron-Cohen S, Fisher SE, Asher JE, et al. Absolute pitch exhibits phenotypic and genetic overlap with synesthesia. Hum Mol Genet. 2013;22:2097–104.

21. Brenton JN, Devries SP, Barton C, Minnich H, Sokol DK. Absolute Pitch in a Four-Year-Old Boy With Autism. Pediatr Neurol. 2008;39:137–8.

22. Heaton P, Davis RE, Happé FGE. Research note: Exceptional absolute pitch perception for spoken words in an able adult with autism. Neuropsychologia. 2008;46:2095–8.

23. Bonnel A, Mottron L, Peretz I, Trudel M, Gallun E, Bonnel AM. Enhanced Pitch Sensitivity in Individuals with Autism: A Signal Detection Analysis. J Cogn Neurosci. 2003;15:226–35.

24. DePape A-MR, Hall GBC, Tillmann B, Trainor LJ. Auditory Processing in High-Functioning Adolescents with Autism Spectrum Disorder. PLoS ONE. 2012;7:e44084.

25. Heaton P, Hudry K, Ludlow A, Hill E. Superior discrimination of speech pitch and its relationship to verbal ability in autism spectrum disorders. Cogn Neuropsychol. 2008;25.

26. Lenhoff HM, Perales O, Hickok G. Absolute Pitch in Williams Syndrome. Music Percept. 2001;18:491–503.

27. Bailey A, Le Couteur A, Gottesman I, Bolton P, Simonoff E, Yuzda E, et al. Autism as a strongly genetic disorder: evidence from a British twin study. Psychol Med. 1995;25:63.

28. Bill BR, Geschwind DH. Genetic advances in autism: heterogeneity and convergence on shared pathways. Curr Opin Genet Dev. 2009;19:271–8.

29. Constantino JN, Zhang Y, Frazier T, Abbacchi AM, Law P. Sibling Recurrence and the Genetic Epidemiology of Autism. Am J Psychiatry. 2010;167:1349–56.

30. Geschwind DH. Genetics of autism spectrum disorders. Trends Cogn Sci. 2011;15:409–16.

31. Persico AM, Napolioni V. Autism genetics. Behav Brain Res. 2013;251:95–112.

32. Bellugi U, Lichtenberger L, Mills D, Galaburda A, Korenberg JR. Bridging cognition, the brain and molecular genetics: evidence from Williams syndrome. Trends Neurosci. 1999;22:197–207.

33. Donnai D, Karmiloff-Smith A. Williams syndrome: From genotype through to the cognitive phenotype. Am J Med Genet. 2000;97:164–71.

34. Meyer-Lindenberg A, Mervis CB, Berman KF. Neural mechanisms in Williams syndrome: a unique window to genetic influences on cognition and behaviour. Nat Rev Neurosci. 2006;7:380–93.

35. Chin CS. The Development of Absolute Pitch: A Theory Concerning the Roles of Music Training at an Early Developmental Age and Individual Cognitive Style. Psychol Music. 2003;31:155–71.

36. Loui P, Li HC, Hohmann A, Schlaug G. Enhanced Cortical Connectivity in Absolute Pitch Musicians: A Model for Local Hyperconnectivity. J Cogn Neurosci. 2011;23:1015–26.

37. Schellenberg EG, Trehub SE. Good Pitch Memory Is Widespread. Psychol Sci. 2003;14:262–6.

38. Russo FA, Windell DL, Cuddy LL. Learning the “Special Note”: Evidence for a Critical Period for Absolute Pitch Acquisition. Music Percept. 2003;21:119–27.

39. Dohn A, Garza-Villarreal EA, Heaton P, Vuust P. Do musicians with perfect pitch have more autism traits than musicians without perfect pitch? An empirical study. PLoS One. 2012;7.

40. Brown WA, Cammuso K, Sachs H, Winklosky B, Mullane J, Bernier R, et al. Autism-Related Language, Personality, and Cognition in People with Absolute Pitch: Results of a Preliminary Study. J Autism Dev Disord. 2003;33:163–7.

41. Jäncke L, Langer N, Hänggi J. Diminished Whole-brain but Enhanced Peri-sylvian Connectivity in Absolute Pitch Musicians. J Cogn Neurosci. 2012;24:1447–61.

42. Courchesne E, Pierce K. Why the frontal cortex in autism might be talking only to itself: local over-connectivity but long-distance disconnection. Curr Opin Neurobiol. 2005;15:225–30.

43. Loui P, Zamm A, Schlaug G. Enhanced functional networks in absolute pitch. NeuroImage. 2012;63:632–40.

44. Belmonte MK. Autism and Abnormal Development of Brain Connectivity. J Neurosci. 2004;24:9228–31.

45. Just MA, Cherkassky VL, Keller TA, Kana RK, Minshew NJ. Functional and Anatomical Cortical Underconnectivity in Autism: Evidence from an fMRI Study of an Executive Function Task and Corpus Callosum Morphometry. Cereb Cortex. 2006;17:951–61.

46. Cherkassky VL, Kana RK, Keller TA, Just MA. Functional connectivity in a baseline resting-state network in autism: NeuroReport. 2006;17:1687–90.

47. Keown CL, Shih P, Nair A, Peterson N, Mulvey ME, Müller R-A. Local Functional Overconnectivity in Posterior Brain Regions Is Associated with Symptom Severity in Autism Spectrum Disorders. Cell Rep. 2013;5:567–72.

48. Lewis JD, Theilmann RJ, Fonov V, Bellec P, Lincoln A, Evans AC, et al. Callosal fiber length and interhemispheric connectivity in adults with autism: Brain overgrowth and underconnectivity. Hum Brain Mapp. 2013;34:1685–95.

49. Murias M, Webb SJ, Greenson J, Dawson G. Resting State Cortical Connectivity Reflected in EEG Coherence in Individuals With Autism. Biol Psychiatry. 2007;62:270–3.

50. Amaral DG, Schumann CM, Nordahl CW. Neuroanatomy of autism. Trends Neurosci. 2008;31:137–45.

51. Mottron L, Dawson M, Soulières I, Hubert B, Burack J. Enhanced Perceptual Functioning in Autism: An Update, and Eight Principles of Autistic Perception. J Autism Dev Disord. 2006;36:27–43.

52. Mottron L, Dawson M, Soulieres I. Enhanced perception in savant syndrome: patterns, structure and creativity. Philos Trans R Soc B Biol Sci. 2009;364:1385–91.

53. Baron-Cohen S. Two new theories of autism: hyper-systemising and assortative mating. Arch Dis Child. 2005;91:2–5.

54. Bargary G, Mitchell KJ. Synaesthesia and cortical connectivity. Trends Neurosci. 2008;31:335–342.

55. Hänggi J, Wotruba D, Jäncke L. Globally altered structural brain network topology in grapheme-color synesthesia. J Neurosci Off J Soc Neurosci. 2011;31:5816–5828.

56. Loui P, Zamm A, Schlaug G. Absolute Pitch and Synesthesia: Two Sides of the Same Coin? Shared and Distinct Neural Substrates of Music Listening. ICMPC Proc Ed Catherine Stevens Al Int Conf Music Percept Cogn. 2012;:618–23.

57. Rouw R, Scholte HS, Colizoli O. Brain areas involved in synaesthesia: A review. J Neuropsychol. 2011;5:214–42.

58. Volberg G, Karmann A, Birkner S, Greenlee MW. Short- and long-range neural synchrony in grapheme-color synesthesia. J Cogn Neurosci. 2013;25:1148–62.

59. Zamm A, Schlaug G, Eagleman DM, Loui P. Pathways to seeing music: Enhanced structural connectivity in colored-music synesthesia. NeuroImage. 2013;74:359–366.

60. Supekar K, Uddin LQ, Khouzam A, Phillips J, Gaillard WD, Kenworthy LE, et al. Brain Hyperconnectivity in Children with Autism and its Links to Social Deficits. Cell Rep. 2013;5:738–47.

61. Mottron L, Bouvet L, Bonnel A, Samson F, Burack JA, Dawson M, et al. Veridical mapping in the development of exceptional autistic abilities. Neurosci Biobehav Rev. 2012;37.

62. Bullmore E, Sporns O. Complex brain networks: graph theoretical analysis of structural and functional systems. Nat Rev Neurosci. 2009;10:186–198.

63. Sporns O. Networks of the Brain. Cambridge, Massachusetts; London, England: MIT Press; 2011.

64. Oldfield RC. The assessment and analysis of handedness: The Edinburgh inventory. Neuropsychologia. 1971;9:97–113.

65. Raven J, Raven JC, Court JH. Manual for Raven’s Progressive Matrices and Vocabulary Tests. Section 3: Standard Progressive Matrices: 2000 Edition, updated 2004. San Antonio: Pearson Assessment; 2004.

66. Oswald WD. Zahlen-Verbindungs-Test (ZVT) - 3., überarbeitete und neu normerte Auflage. 3rd edition. Göttingen: Hogrefe; 2016.

67. Gordon EE. Manual for the advanced measures of music audiation. GIA Publications; 1989.

68. Müllensiefen D, Gingras B, Musil J, Stewart L. The Musicality of Non-Musicians: An Index for Assessing Musical Sophistication in the General Population. PLOS ONE. 2014;9:e89642.

69. Dohn A, Garza-Villarreal EA, Ribe LR, Wallentin M, Vuust P. Musical Activity Tunes Up Absolute Pitch Ability. Music Percept Interdiscip J. 2014;31:359–71.

70. Baron-Cohen S, Wheelwright S, Skinner R, Martin J, Clubley E. The Autism-Spectrum Quotient (AQ): Evidence from Asperger Syndrome/High-Functioning Autism, Malesand Females, Scientists and Mathematicians. J Autism Dev Disord. 2001;31:5–17.

71. Peirce JW. PsychoPy—Psychophysics software in Python. J Neurosci Methods. 2007;162:8–13.

72. Delorme A, Makeig S. EEGLAB: an open source toolbox for analysis of single-trial EEG dynamics including independent component analysis. J Neurosci Methods. 2004;134:9–21.

73. Oostenveld R, Fries P, Maris E, Schoffelen J-M. FieldTrip: Open Source Software for Advanced Analysis of MEG, EEG, and Invasive Electrophysiological Data. Comput Intell Neurosci. 2011;2011:1–9.

74. Perrin F, Pernier J, Bertrand O, Echallier JF. Spherical splines for scalp potential and current density mapping. Electroencephalogr Clin Neurophysiol. 1989;72:184–7.

75. Vinck M, Battaglia FP, Womelsdorf T, Pennartz C. Improved measures of phase-coupling between spikes and the Local Field Potential. J Comput Neurosci. 2012;33:53–75.

76. Plonsey R, Heppner DB. Considerations of quasi-stationarity in electrophysiological systems. Bull Math Biophys. 1967;29:657–64.

77. Stinstra JG, Peters MJ. The volume conductor may act as a temporal filter on the ECG and EEG. Med Biol Eng Comput. 1998;36:711–6.

78. Nunez PL, Srinivasan R, Westdorp AF, Wijesinghe RS, Tucker DM, Silberstein RB, et al. EEG coherency. Electroencephalogr Clin Neurophysiol. 1997;103:499–515.

79. Stam CJ, Nolte G, Daffertshofer A. Phase lag index: Assessment of functional connectivity from multi channel EEG and MEG with diminished bias from common sources. Hum Brain Mapp. 2007;28:1178–93.

80. Cohen MX. Analyzing Neural Time Series Data. Theory and Practice. Cambridge, Massachusetts; London, England: MIT Press; 2014.

81. Mormann F, Lehnertz K, David P, E. Elger C. Mean phase coherence as a measure for phase synchronization and its application to the EEG of epilepsy patients. Phys Nonlinear Phenom. 2000;144:358–69.

82. Stam C, Jones B, Nolte G, Breakspear M, Scheltens P. Small-World Networks and Functional Connectivity in Alzheimer’s Disease. Cereb Cortex. 2006;17:92–9.

83. Ortiz E, Stingl K, Münßinger J, Braun C, Preissl H, Belardinelli P. Weighted Phase Lag Index and Graph Analysis: Preliminary Investigation of Functional Connectivity during Resting State in Children. Comput Math Methods Med. 2012;2012:1–8.

84. Hardmeier M, Hatz F, Bousleiman H, Schindler C, Stam CJ, Fuhr P. Reproducibility of Functional Connectivity and Graph Measures Based on the Phase Lag Index (PLI) and Weighted Phase Lag Index (wPLI) Derived from High Resolution EEG. PLoS ONE. 2014;9:e108648.

85. Rubinov M, Sporns O. Complex network measures of brain connectivity: uses and interpretations. NeuroImage. 2010;52:1059–69.

86. Langer N, Pedroni A, Gianotti LRR, Hänggi J, Knoch D, Jäncke L. Functional brain network efficiency predicts intelligence. Hum Brain Mapp. 2012;33:1393–406.

87. de Haan W, Pijnenburg YA, Strijers RL, van der Made Y, van der Flier WM, Scheltens P, et al. Functional neural network analysis in frontotemporal dementia and Alzheimer’s disease using EEG and graph theory. BMC Neurosci. 2009;10:101.

88. Iturria-Medina Y, Sotero RC, Canales-Rodríguez EJ, Alemán-Gómez Y, Melie-García L. Studying the human brain anatomical network via diffusion-weighted MRI and Graph Theory. NeuroImage. 2008;40:1064–76.

89. Iturria-Medina Y, Canales-Rodríguez EJ, Melie-García L, Valdés-Hernández PA, Martínez-Montes E, Alemán-Gómez Y, et al. Characterizing brain anatomical connections using diffusion weighted MRI and graph theory. NeuroImage. 2007;36:645–60.

90. Zhou Y, Yu F, Duong T. Multiparametric MRI Characterization and Prediction in Autism Spectrum Disorder Using Graph Theory and Machine Learning. PLoS ONE. 2014;9:e90405.

91. van Wijk, Bernadette C. M., Stam CJ, Daffertshofer A, Sporns O. Comparing Brain Networks of Different Size and Connectivity Density Using Graph Theory. PloS One. 2010;5:e13701.

92. Stam CJ, Reijneveld JC. Graph theoretical analysis of complex networks in the brain. Nonlinear Biomed Phys. 2007;1:3.

93. Sporns O, Zwi JD. The Small World of the Cerebral Cortex. Neuroinformatics. 2004;2:145–62.

94. van Straaten, E. C. W., Stam CJ. Structure out of chaos: functional brain network analysis with EEG, MEG, and functional MRI. Eur Neuropsychopharmacol J Eur Coll Neuropsychopharmacol. 2013;23:7–18.

95. Latora V, Marchiori M. Efficient Behavior of Small-World Networks. Phys Rev Lett. 2001;87. doi:10.1103/PhysRevLett.87.198701.

96. Cohen JR, D’Esposito M. The Segregation and Integration of Distinct Brain Networks and Their Relationship to Cognition. J Neurosci. 2016;36:12083–94.

97. Watts DJ, Strogatz SH. Collective dynamics of “small-world” networks. Nature. 1998;:441–2.

98. Bullmore E, Sporns O. The economy of brain network organization. Nat Rev Neurosci. 2012;13:336–349.

99. Achard S, Bullmore E. Efficiency and cost of economical brain functional networks. PLoS Comput Biol. 2007;3:e17.

100. Senkowski D, Schneider TR, Foxe JJ, Engel AK. Crossmodal binding through neural coherence: implications for multisensory processing. Trends Neurosci. 2008;31:401–9.

101. Xia M, Wang J, He Y. BrainNet Viewer: A Network Visualization Tool for Human Brain Connectomics. PLoS ONE. 2013;8:e68910.

102. Koessler L, Maillard L, Benhadid A, Vignal JP, Felblinger J, Vespignani H, et al. Automated cortical projection of EEG sensors: Anatomical correlation via the international 10–10 system. NeuroImage. 2009;46:64–72.

103. Peters JM, Taquet M, Vega C, Jeste SS, Fernández IS, Tan J, et al. Brain functional networks in syndromic and non-syndromic autism: a graph theoretical study of EEG connectivity. BMC Med. 2013;11. doi:10.1186/1741-7015-11-54.

104. Cantero JL, Atienza M, Madsen JR, Stickgold R. Gamma EEG dynamics in neocortex and hippocampus during human wakefulness and sleep. NeuroImage. 2004;22:1271–80.

105. Bragin A, Jando G, Nadasdy Z, Hetke J, Wise K, Buzsaki G. Gamma (40-100 Hz) oscillation in the hippocampus of the behaving rat. J Neurosci. 1995;15:47–60.

106. Miltner WHR, Braun C, Arnold M, Witte H, Taub E. Coherence of gamma-band EEG activity as a basis for associative learning. Nature. 1999;397:434–6.

107. Herrmann CS, Fründ I, Lenz D. Human gamma-band activity: A review on cognitive and behavioral correlates and network models. Neurosci Biobehav Rev. 2010;34:981–92.

108. Elmer S, Rogenmoser L, Kühnis J, Jäncke L. Bridging the Gap between Perceptual and Cognitive Perspectives on Absolute Pitch. J Neurosci. 2015;35:366–71.

109. Zatorre RJ, Beckett C. Multiple coding strategies in the retention of musical tones by possessors of absolute pitch. Mem Cognit. 1989;17:582–9.

110. Schulze K, Gaab N, Schlaug G. Perceiving pitch absolutely: Comparing absolute and relative pitch possessors in a pitch memory task. BMC Neurosci. 2009;10:106.

111. Levitin DJ. Absolute memory for musical pitch: Evidence from the production of learned melodies. Percept Psychophys. 1994;56:414–23.

112. Bermudez P, Zatorre RJ. The absolute pitch mind continues to reveal itself. J Biol. 2009;8:75.

113. Bhattacharya J, Petsche H, Pereda E. Long-Range Synchrony in the Gamma Band: Role in Music Perception. J Neurosci. 2001;21:6329–6337.

114. Bhattacharya J, Petsche H. Musicians and the gamma band: A secret affair? NeuroReport. 2001;12:371–374.

115. Sun L, Grutzner C, Bolte S, Wibral M, Tozman T, Schlitt S, et al. Impaired Gamma-Band Activity during Perceptual Organization in Adults with Autism Spectrum Disorders: Evidence for Dysfunctional Network Activity in Frontal-Posterior Cortices. J Neurosci. 2012;32:9563–73.

116. Brown C, Gruber T, Boucher J, Rippon G, Brock J. Gamma Abnormalities During Perception of Illusory Figures in Autism. Cortex. 2005;41:364–76.

117. Herrmann C, Demiralp T. Human EEG gamma oscillations in neuropsychiatric disorders. Clin Neurophysiol. 2005;116:2719–33.

118. Uhlhaas PJ, Singer W. Neural synchrony in brain disorders: relevance for cognitive dysfunctions and pathophysiology. Neuron. 2006;52:155–68.

119. Brock J, Brown CC, Boucher J, Rippon G. The temporal binding deficit hypothesis of autism. Dev Psychopathol. 2002;14. doi:10.1017/S0954579402002018.

120. Grice SJ, Spratling MW, Karmiloff-Smith A, Halit H, Csibra G, de Haan M, et al. Disordered visual processing and oscillatory brain activity in autism and Williams Syndrome. NeuroReport. 2001;12:2697.

121. Brosnan MJ, Scott FJ, Fox S, Pye J. Gestalt processing in autism: failure to process perceptual relationships and the implications for contextual understanding. J Child Psychol Psychiatry. 2004;45:459–69.

122. Happé F, Frith U. The weak coherence account: detail-focused cognitive style in autism spectrum disorders. J Autism Dev Disord. 2006;36.

123. Dawson G, Webb S, Schellenberg GD, Dager S, Friedman S, Aylward E, et al. Defining the broader phenotype of autism: Genetic, brain, and behavioral perspectives. Dev Psychopathol. 2002;14. doi:10.1017/S0954579402003103.

124. Samson F, Mottron L, Soulières I, Zeffiro TA. Enhanced visual functioning in autism: an ALE meta-analysis. Hum Brain Mapp. 2012;33.

125. Happé F. Autism: cognitive deficit or cognitive style? Trends Cogn Sci. 1999;3.

126. Happé F, Frith U. The neuropsychology of autism. Brain. 1996;119.

127. Frith U. Autism: Explaining the Enigma. Oxford, UK: Blackwell; 1989.

128. Costa-Giomi E, Gilmour R, Siddell J, Lefebvre E. Absolute Pitch, Early Musical Instruction, and Spatial Abilities. Ann N Y Acad Sci. 2006;930:394–6.

129. Gervain J, Vines BW, Chen LM, Seo RJ, Hensch TK, Werker JF, et al. Valproate reopens critical-period learning of absolute pitch. Front Syst Neurosci. 2013;7.doi:10.3389/fnsys.2013.00102.

130. Christensen J, Grønborg TK, Sørensen MJ, Schendel D, Parner ET, Pedersen LH, et al. Prenatal Valproate Exposure and Risk of Autism Spectrum Disorders and Childhood Autism. JAMA. 2013;309:1696.

131. Williams G, King J, Cunningham M, Stephan M, Kerr B, Hersh JH. Fetal valproate syndrome and autism: additional evidence of an association. Dev Med Child Neurol. 2001;43:202–6.

132. Dufour-Rainfray D, Vourc’h P, Le Guisquet A-M, Garreau L, Ternant D, Bodard S, et al. Behavior and serotonergic disorders in rats exposed prenatally to valproate: A model for autism. Neurosci Lett. 2010;470:55–9.

133. Wagner GC, Reuhl KR, Cheh M, McRae P, Halladay AK. A New Neurobehavioral Model of Autism in Mice: Pre- and Postnatal Exposure to Sodium Valproate. J Autism Dev Disord. 2006;36:779–93.

134. Moseley RL, Ypma RJF, Holt RJ, Floris D, Chura LR, Spencer MD, et al. Whole-brain functional hypoconnectivity as an endophenotype of autism in adolescents. NeuroImage Clin. 2015;9:140–52.

135. Pellicano E, Gibson L, Maybery M, Durkin K, Badcock DR. Abnormal global processing along the dorsal visual pathway in autism: a possible mechanism for weak visuospatial coherence? Neuropsychologia. 2005;43:1044–53.

136. Lo Y-C, Soong W-T, Gau SS-F, Wu Y-Y, Lai M-C, Yeh F-C, et al. The loss of asymmetry and reduced interhemispheric connectivity in adolescents with autism: A study using diffusion spectrum imaging tractography. Psychiatry Res Neuroimaging. 2011;192:60–6.

137. Dohn A, Garza-Villarreal EA, Chakravarty MM, Hansen M, Lerch JP, Vuust P. Gray- and White-Matter Anatomy of Absolute Pitch Possessors. Cereb Cortex. 2015;25:1379–88.

138. Tsiaras V, Simos PG, Rezaie R, Sheth BR, Garyfallidis E, Castillo EM, et al. Extracting biomarkers of autism from MEG resting-state functional connectivity networks. Comput Biol Med. 2011;41:1166–77.

139. Sihvonen AJ, Ripollés P, Särkämö T, Leo V, Rodríguez-Fornells A, Saunavaara J, et al. Tracting the neural basis of music: Deficient structural connectivity underlying acquired amusia. Cortex. 2017;97:255–73.

140. Sihvonen AJ, Ripolles P, Leo V, Rodriguez-Fornells A, Soinila S, Sarkamo T. Neural Basis of Acquired Amusia and Its Recovery after Stroke. J Neurosci. 2016;36:8872–81.

141. Schlaug G, Jäncke L, Huang Y, Staiger JF, Steinmetz H. Increased corpus callosum size in musicians. Neuropsychol Dev Stud Corpus Callosum. 1995;33:1047–55.

142. Bengtsson SL, Nagy Z, Skare S, Forsman L, Forssberg H, Ullén F. Extensive piano practicing has regionally specific effects on white matter development. Nat Neurosci. 2005;8:1148–50.

143. Burunat I, Brattico E, Puoliväli T, Ristaniemi T, Sams M, Toiviainen P. Action in Perception: Prominent Visuo-Motor Functional Symmetry in Musicians during Music Listening. PLOS ONE. 2015;10:e0138238.

144. Schmithorst VJ, Wilke M. Differences in white matter architecture between musicians and non-musicians: a diffusion tensor imaging study. Neurosci Lett. 2002;321:57–60.

145. Elmer S, Hänggi J, Jäncke L. Interhemispheric transcallosal connectivity between the left and right planum temporale predicts musicianship, performance in temporal speech processing, and functional specialization. Brain Struct Funct. 2014.

146. Sihvonen AJ, Särkämö T, Ripollés P, Leo V, Saunavaara J, Parkkola R, et al. Functional neural changes associated with acquired amusia across different stages of recovery after stroke. Sci Rep. 2017;7:11390.

147. Sota S, Hatada S, Honjyo K, Takatsuka T, Honer WG, Morinobu S, et al. Musical disability in children with autism spectrum disorder. Psychiatry Res. 2018;267:354–9.

148. von Stein A, Sarnthein J. Different frequencies for different scales of cortical integration: from local gamma to long range alpha/theta synchronization. Int J Psychophysiol. 2000;38:301–13.

149. Wass S. Distortions and disconnections: disrupted brain connectivity in autism. Brain Cogn. 2011;75.

150. Heaton P. Pitch memory, labelling and disembedding in autism. J Child Psychol Psychiatry. 2003;44:543–51.

